# Sleep deprivation negatively impacts reproductive output in *Drosophila melanogaster*

**DOI:** 10.1101/158071

**Authors:** Sheetal Potdar, Danita K. Daniel, Femi A. Thomas, Shraddha Lall, Vasu Sheeba

## Abstract

Most animals sleep or exhibit a sleep-like state, yet the adaptive significance of this phenomenon remains unclear. Although reproductive deficits are associated with lifestyle induced sleep deficiencies, how sleep loss affects reproductive physiology is poorly understood, even in model organisms. We aimed to bridge this mechanistic gap by impairing sleep in female fruit flies and testing its effect on egg output. We find that sleep deprivation by feeding caffeine or by mechanical perturbation results in decreased egg output. Transient activation of wake-promoting dopaminergic neurons decreases egg output in addition to sleep levels, thus demonstrating a direct negative impact of sleep deficit on reproductive output. Similarly, loss-of-function mutation in dopamine transporter *fumin* (*fmn*)leads to both significant sleep loss and lowered fecundity. This demonstration of a direct relationship between sleep and reproductive fitness indicates a strong driving force for the evolution of sleep.

## Introduction

Almost all animals show activity/rest cycles in response to daily solar cycles of light, temperature and other environmental cues. The rest phase of sleep is remarkably ubiquitous in animals suggesting that sleep is important. While we humans spend a third of our lives sleeping, we do not know *why* sleep is indispensable. Several studies link sleep levels to cognition, mood and emotional states (Krause et al., 2017), as well as physiological health in humans (Mahoney, 2010). When rats are chronically deprived of sleep there are detrimental effects on longevity (Rechtschaffen, Gilliland, Bergmann, & Winter, 1983), skin condition (Everson, Bergmann, & Rechtschaffen, 1989) and body weight (Everson & Szabo, 2011) accompanied by physiological changes in internal organs (Everson & Szabo, 2009). Thus, sleep positively influences many organ systems in addition to the nervous system.

The genetically tractable model organism *Drosophila melanogaster* exhibits several characteristics of mammalian sleep - increased arousal threshold, site-specificity, regulation by homeostatic and circadian clock mechanisms and even sleep-specific electrophysiological signatures (Hendricks et al., 2000; Nitz, van Swinderen, Tononi, & Greenspan, 2002; Shaw, Cirelli, Greenspan, & Tononi, 2000; van Alphen, Yap, Kirszenblat, Kottler, & van Swinderen, 2013). Sleep deprivation in flies results in deleterious effects similar to those seen in mammals. Mechanically depriving flies of sleep decreases their lifespan (Seugnet et al., 2009; Shaw, Tononi, Greenspan, & Robinson, 2002) and short-sleeping mutants of the Shaker potassium channel have reduced lifespan (Bushey, Hughes, Tononi, & Cirelli, 2010; Cirelli et al., 2005). However, lifespan by itself is an insufficient indicator of overall fitness of an organism as it can be radically influenced by reproductive output (Sheeba, Sharma, Shubha, Chandrashekaran, & Joshi, 2000). Since reproductive success is a strong evolutionary driving force, we focused on possible mechanistic links between sleep and reproductive output.

In humans, infertility is often associated with sleep disturbances; however, the complexity of the reproductive system and sleep characteristics in humans makes the analysis of sleep disruption affecting reproductive processes difficult (Kloss, Perlis, Zamzow, Culnan, & Gracia, 2015). Shift workers and women who experience frequent jet lag conditions report sleep disturbances and abnormal menstrual cycles and are at a higher risk of developing pregnancy-related complications (Mahoney, 2010). Chronic sleep deprivation in rats increases spontaneous ejaculations (Andersen & Tufik, 2002) and reduces the number of live sperm (Alvarenga, Hirotsu, Mazaro-Costa, Tufik, & Andersen, 2015). In mice subjected to light protocols mimicking jet lag and circadian misalignment, reproductive success is hampered (Summa, Vitaterna, & Turek, 2012). Circadian clock mutants with defective timing and consolidation of sleep also have reduced reproductive output in flies (Beaver et al., 2002) and mice (Loh et al., 2014). Sleep deprivation alters aggressive behaviour in flies and hampers the chances of mating (Kayser, Mainwaring, Yue, & Sehgal, 2015). Most studies show that sleep and reproductive output are associated with one another, without testing the direct effects of sleep on reproductive success. Here, we address this question by impairing sleep in female fruit flies and testing its effect on reproductive output. We find that feeding flies with caffeine or depriving them of sleep by mechanical perturbation, or by decreasing sleep by genetic activation of wake-promoting dopamine neurons all result in decreased egg output. Decreased sleep is associated with decreased egg output for all manipulations. Thus, our study establishes a model system to study the mechanisms underlying relationships between sleep and reproductive processes that underlie fitness.

## Results

### Effect of sleep deprivation on egg output of inbred *w*^1118^ flies

To assess the impact of sleep deprivation upon reproductive output, we first used caffeine to deprive female flies of sleep. Flies were given caffeinated food during the day only (D_caf_), or during the night only (N_caf_) or standard cornmeal food during both day and night that acted as controls (Ctrl). To estimate the appropriate concentration of caffeine for our egg output assay, we quantified the amount of sleep loss in flies with two concentrations (0.5 and 1 mg/ml) based on previous studies (Andretic, Kim, Jones, Han, & Greenspan, 2008; Wu et al., 2009) and our pilot experiments. Flies that were fed with food containing 0.5 mg/ml caffeine only during the day (D_caf_) tend to exhibit less sleep during the day as compared to their own baseline (BS) as well as compared to control flies during caffeine (CAF) days (Fig 1A, BS and CAF), although this reduction was not statistically significant (Fig 1B, day). However, these flies showed a rebound increase in daytime sleep upon removal from caffeinated food (Fig 1A, RC) which was significantly higher than daytime sleep during BS and CAF (Fig 1B, day). Similarly, when flies were fed with food containing 0.5 mg/ml caffeine only during the night (N_caf_, Fig 1A-B,), their night sleep was significantly reduced as compared to their own BS days as well as control flies during CAF days (Fig 1A, BS and CAF; Fig 1B, night). These data show that caffeine has an immediate effect on sleep – D_caf_ flies show reduced daytime sleep while N_caf_ flies show reduced night sleep. We found similar trends of reduced daytime sleep of D_caf_ and reduced night sleep of N_caf_ with respect to BS when flies were fed with food containing 1 mg/ml caffeine (Supplementary Fig 1). Importantly, 0.5 mg/ml is more efficient in decreasing sleep levels (53% day and 49% night sleep loss) as compared to 1.0 mg/ml of caffeine (38 % day and 4 % night sleep loss, Fig 1B’). This may be due to reduced food intake with increasing caffeine content, which could in turn result in lesser extent of sleep loss.

**Figure 1.**
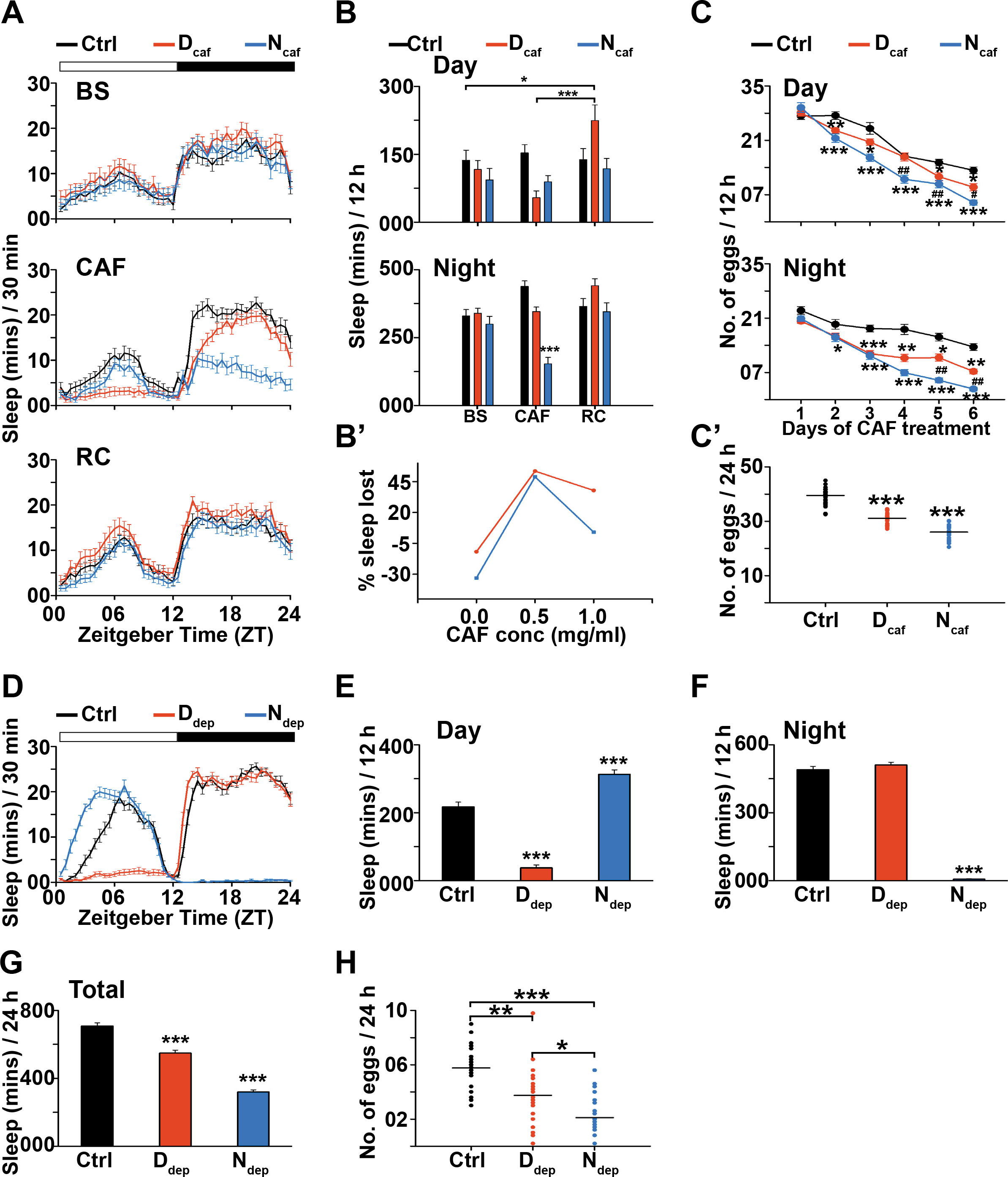
Sleep deprivation by caffeine and mechanical disturbance of *w*^1118^ flies results in decrease of egg output. (A) Sleep in minutes for every half hour over a period of 24 h is shown for w^1118^ flies fed with standard food (Ctrl, *n* = 28), flies fed with 0.5 mg/ml caffeine only during the day (D_caf_, *n* = 25) and only during the night (N_caf_, *n* = 24) averaged across two baseline (BS), three caffeine feeding (CAF) and two recovery (RC) days. Horizontal white and black bars on top represent day and night respectively. (B) Daytime (top) and night (bottom) sleep of control, Dcaf and Ncaf flies are compared across BS, CAF and RC days. Dcaf flies show significant increase in daytime sleep during RC days as compared to that during BS and CAF days. N_caf_ flies show significantly lower levels of night sleep during CAF days as compared to that during BS and RC days, as well as night sleep of controls during CAF days (two-way ANOVA with treatment and days as fixed factors followed by post-hoc Tukey’s HSD test). (B’) Percentage total sleep loss during CAF days with respect to BS days plotted as function of caffeine concentration shows that sleep loss is higher for caffeine concentration of 0.5 mg/ml during both day and night as compared to a concentration of 1.0 mg/ml. (C) Number of eggs laid by control (n = 25), D_caf_ (n = 24) and N_caf_ (n = 25) flies both during day and night over a period of six days of caffeine (0.5 mg/ml) treatment. * denotes significant differences between either Dcaf or Ncaf with control flies, while # indicates significant differences between D_caf_ and N_ca_f flies (Kruskal-Wallis test). (C’) Total number of eggs laid averaged across six days of caffeine treatment. D_caf_ flies laid significantly lesser number of eggs as compared to control flies, while N_caf_ flies lay significantly lower number of eggs as compared to both control and D_caf_ flies (one-way ANOVA with treatment as fixed factor followed by post-hoc Tukey’s HSD test). The experiment was repeated with similar results (data not shown). (D) Sleep in minutes for every half hour over a period of 24 h averaged across 5 days is shown for control w^1118^ flies (Ctrl, *n* = 26), flies receiving mechanical disturbance only during the day (D_dep_, *n* = 28) and only during the night (N_dep_, *n* = 27). (E) Daytime sleep of D_dep_ flies significantly reduced as compared to Ctrl and N_dep_, whereas that of N_dep_ flies significantly higher than that of Ctrl and D_dep_. (F) Night sleep of N_dep_ flies significantly lower than Ctrl and D_dep_ flies. (G) Total sleep of D_dep_ flies is significantly lower than Ctrl and that of Ndep flies is significantly lower than Ctrl and Ddep flies (one way ANOVA with treatment as fixed factor followed by post-hoc Tukey’s HSD test for E, F and G). (H) Total number of eggs laid by Ctrl, D_dep_ and N_dep_ flies averaged across 5 days. D_dep_ flies show significant reduction in number of eggs laid as compared to Ctrl; N_dep_ flies laid even lower number of eggs significantly reduced as compared to both Ctrl and D_dep_ flies (Kruskal-Wallis test). * *p* < 0.05, ** *p* < 0.005, *** *p* < 0.0005. Error bars are SEM.

Since providing flies with food containing 0.5 mg/ml caffeine during day or night leads to about 50 % reduction in both daytime and night sleep respectively, we next determined how this affects their reproductive output. We subjected 5-day old female flies (mated for one day prior to the start of the experiment) to caffeine treatment only during the day (D_caf_) or only during the night (N_caf_). We found that both Dcaf and Ncaf flies laid lesser number of eggs as compared to the control flies both during the day as well as night (Fig 1C). N_caf_ flies laid lesser number of eggs as compared to D_caf_ flies also, which was statistically significant on the later days of the treatment (Fig 1C). When we compared the total number of eggs averaged over the 6 days of treatment, D_caf_ flies laid significantly lesser number of eggs as compared to control flies, and Ncaf flies laid significantly lesser number of eggs as compared to both control and D_caf_ flies (Fig 1C’).

Since it is likely that flies fed with caffeine laid fewer eggs simply because oviposition was inhibited by food containing caffeine, we carried out an oviposition preference assay, where flies were allowed to lay eggs on a petri plate, with half the plate containing standard food and the other half containing 0.5 mg/ml caffeinated food. We found that flies laid almost equal number of eggs on both halves, suggesting that for food containing caffeine at a concentration of 0.5 mg/ml, flies do not have any ovipositional avoidance (Preference Index _caf_ = 0.49 ± 0.11, chi-square test, *χ*^2^ = 0.049, *p* = 0.82). Overall, these results suggest that caffeine decreases egg output and flies that lose night sleep tend to lay lesser number of eggs than flies that lose daytime sleep.

To confirm the effect of sleep loss in egg output we used a completely different sleep deprivation method. We substituted caffeine with a vortexer-based mechanical perturbation protocol. Three sets of flies received either of the following treatments - exposure to mechanical disturbance only during day (D_dep_), or only during night (N_dep_) or control (Ctrl) condition with no mechanical perturbation. For the same sets of flies, we obtained both sleep levels and egg counts by transferring flies to fresh tubes every 12 hours for five days. As expected, mechanical disturbance during day reduced daytime sleep and that during night reduced night sleep drastically (Fig 1D-F,). However, only N_dep_ flies recovered this lost night sleep during the subsequent days (Fig 1E) whereas D_dep_ flies did not recover the lost daytime sleep during subsequent nights (Fig 1F). Nevertheless, N_dep_ flies lost greater amount of overall sleep as compared to D_dep_ flies (Fig 1G). Importantly, the average egg output in both D_dep_ and N_dep_ flies was significantly lowered as compared to the control flies (Fig 1H). Furthermore, N_dep_ flies, which on average lost more sleep, also laid significantly lesser number of eggs as compared to D_dep_ flies (Fig 1G-H,). Thus, these results along with similar results obtained with sleep deprivation using caffeine suggest that sleep loss results in reduction in egg output and that sleep loss during the night has a greater detrimental effect on egg output.

### Effect of sleep deprivation on reproductive fitness of outbred flies

We used a strain of *w*^1118^ flies which has been maintained in our laboratory for several years and is likely to harbour loci that have been fixed for certain traits which may have resulted in the above phenotype by chance. Given that reproductive output is a major Darwinian fitness trait, we asked how sleep loss might affect reproductive output in a large, random mating and therefore outbred population of flies which is unlikely to have suffered from similar genetic bottlenecks (CCM) (Gogna, Singh, Sheeba, & Dorai, 2015). We subjected flies to three different concentrations of caffeine (0.5, 1.0 and 1.5 mg/ml) either only during day or only during night and found that none of the D_caf_ flies lost daytime sleep, whereas all the N_caf_ flies lost similar amounts of night sleep (Fig 2A-B,). However, D_caf_ (1.5 mg/ml) flies laid significantly lower number of eggs than the control flies, suggesting that caffeine can affect egg output even without its effect on daytime sleep (Fig 2C). Moreover, N_caf_ flies receiving 0.5 mg/ml and 1.5 mg/ml caffeine also showed reduced egg output as compared to control flies (Fig 2C). These results point toward a direct effect of caffeine on egg output independent of its effect on sleep as well as an indirect effect on egg output through sleep loss. To probe this further, we increased caffeine concentration and found that even higher caffeine concentrations of 4.0 mg/ml fed during the day did not affect daytime sleep (Supplementary Fig 2A-B,S and CAF, 2B, day), however, when fed during the night, decreased night sleep (Supplementary Fig 2B, night). With respect to egg output, we found that the total number of eggs laid by D_caf_ and N_caf_ flies was significantly lower than that of the control flies, however, the number of eggs laid by D_caf_ and N_caf_ flies were not statistically different from each other (Supplementary Fig 2C) similar to what was found for lower concentrations of caffeine. Caffeine treatment does not affect the viability of the eggs laid as seen from egg-to-adult survivorship of eggs laid by D_caf_, N_caf_ (0.5 mg/ml) and Crtl flies (data not shown). Taken together, these results suggest that caffeine treatment may affect the reproductive fitness directly or indirectly through sleep loss.

**Figure 2.**
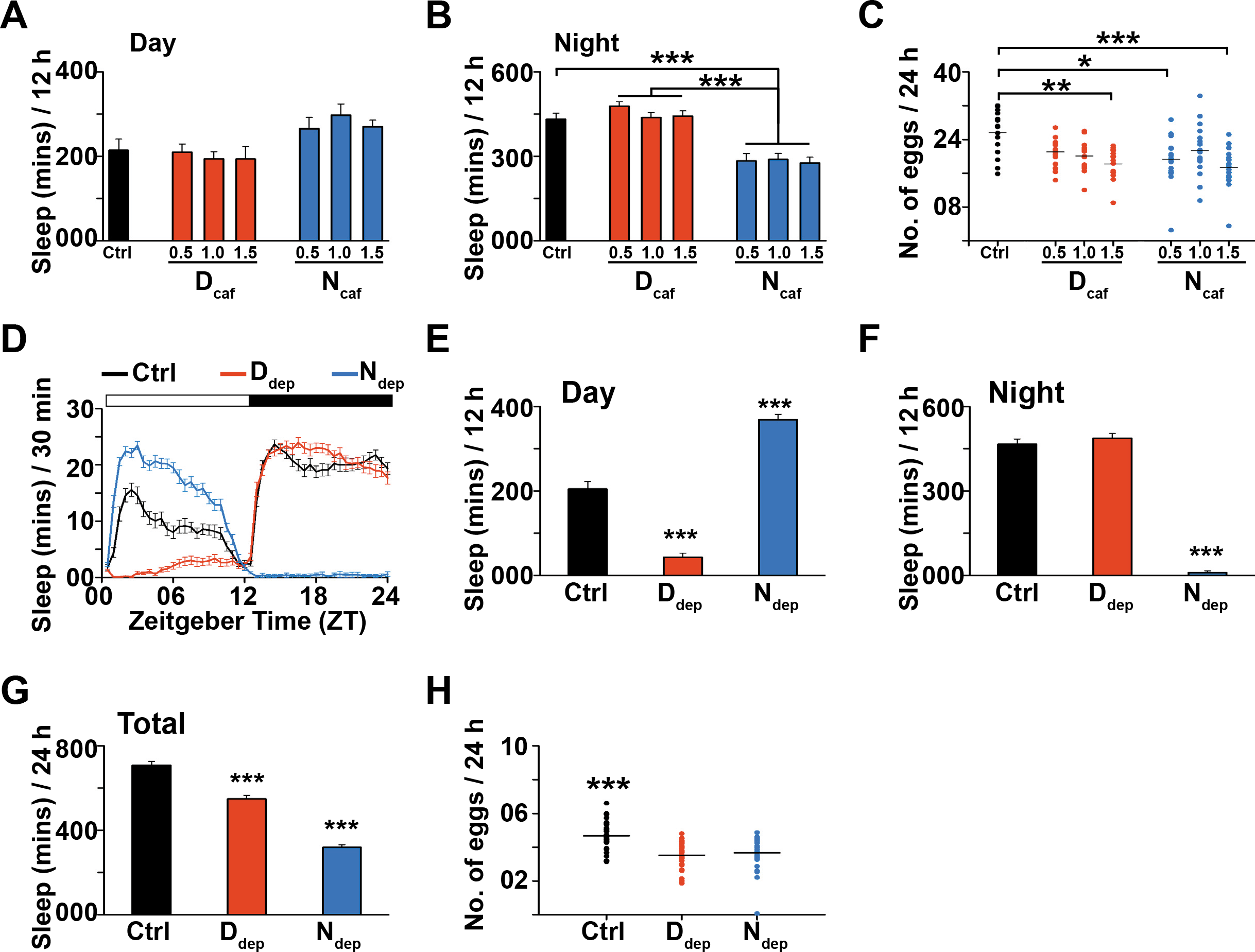
Sleep deprivation by caffeine and mechanical disturbance of outbred CCM flies results in egg output reduction. (A) Daytime and (B) night sleep of flies of outbred CCM population fed with standard food, or caffeine food of different concentrations (0.5, 1.0 and 1.5 mg/ml) either only during day (D_caf_) or only during night (N_caf_). Daytime sleep of flies receiving all the treatments is similar, while night sleep of Ncaf flies of all caffeine concentrations is significantly reduced as compared to control and D_caf_ flies of all caffeine concentrations (one-way ANOVA with treatment as fixed factor followed by post-hoc Tukey’s HSD test). *n* > 21 for all treatments. (C) Total number of eggs laid averaged across 6 days by D_caf_-1.5 (n = 13), N_caf_-0.5 (n = 19) and N_caf_-1.5 (n = 17) flies are significantly reduced as compared to the control (n = 16) flies. D_caf_ and N_caf_ flies of any caffeine concentration do not differ in the total number of eggs laid from each other. D_caf_-0.5 (n = 17), D_caf_-1.0 (n = 17) and N_caf_-1.0 (n = 18) do not differ from the control flies in the number of eggs laid (Kruskal-Wallis test). (D) Sleep in minutes for every half hour over a period of 24 h averaged across five days is shown for control (n = 28) flies of outbred CCM population, flies mechanically disturbed during the day (D_dep_, *n* = 30) and during the night (N_dep_, *n* = 31). (E) During the day, D_dep_ flies sleep significantly lower than both control and Ndep flies due to mechanical disturbance, Ndep flies sleep significantly higher than control and D_dep_ flies indicating sleep rebound due to sleep deprivation during the previous night. (F) During the night, Ndep flies sleep significantly lower than the control and Ddep flies due to mechanical perturbation. (G) Total sleep averaged across 5 days of D_dep_ flies is significantly lower than control flies, whereas that of Ndep is significantly lower than both control and Ddep flies (one-way ANOVA with treatment as fixed factor followed by post-hoc Tukey’s HSD test for E, F and G). (H) Total number of eggs laid averaged across five days by both D_dep_ and N_dep_ flies is significantly lower as compared to control flies (Kuskal-Wallis test). All other details as in Figure 1. A similar experiment with higher levels of deprivation yielded similar results (data not shown).

We next subjected the CCM flies to sleep deprivation protocol using mechanical perturbation either during the day only (D_dep_) or during the night only (N_dep_). Expectedly, D_dep_ flies lost day sleep and N_dep_ flies lost night sleep which they could recover during subsequent days (Fig 2D-F,). Nevertheless N_dep_ flies lost overall greater amount of sleep as compared to D_dep_ flies (Fig 2G). Again, as in the case of caffeine fed outbred flies, with mechanical disturbance also we found that there is a reduction in egg output in D_dep_ and N_dep_ flies as compared to control flies, although there was no difference in egg output between flies experiencing day vs. night sleep disturbance (Fig 2H). However, in yet another assay with mechanically sleep deprived flies, egg output of N_dep_ flies averaged across three days *after* the deprivation protocol was still significantly reduced, while that of Ddep flies was comparable to control flies (Supplementary Fig 3). Therefore, with both caffeine and mechanical disturbance, the resultant sleep deprivation contributed in part to the decrease in egg output of outbred flies. Furthermore, as seen in inbred flies, night sleep loss had greater impact on egg output as compared to daytime sleep loss, though this difference was less discernible and the effect much more subtle in outbred flies.

**Figure 3.**
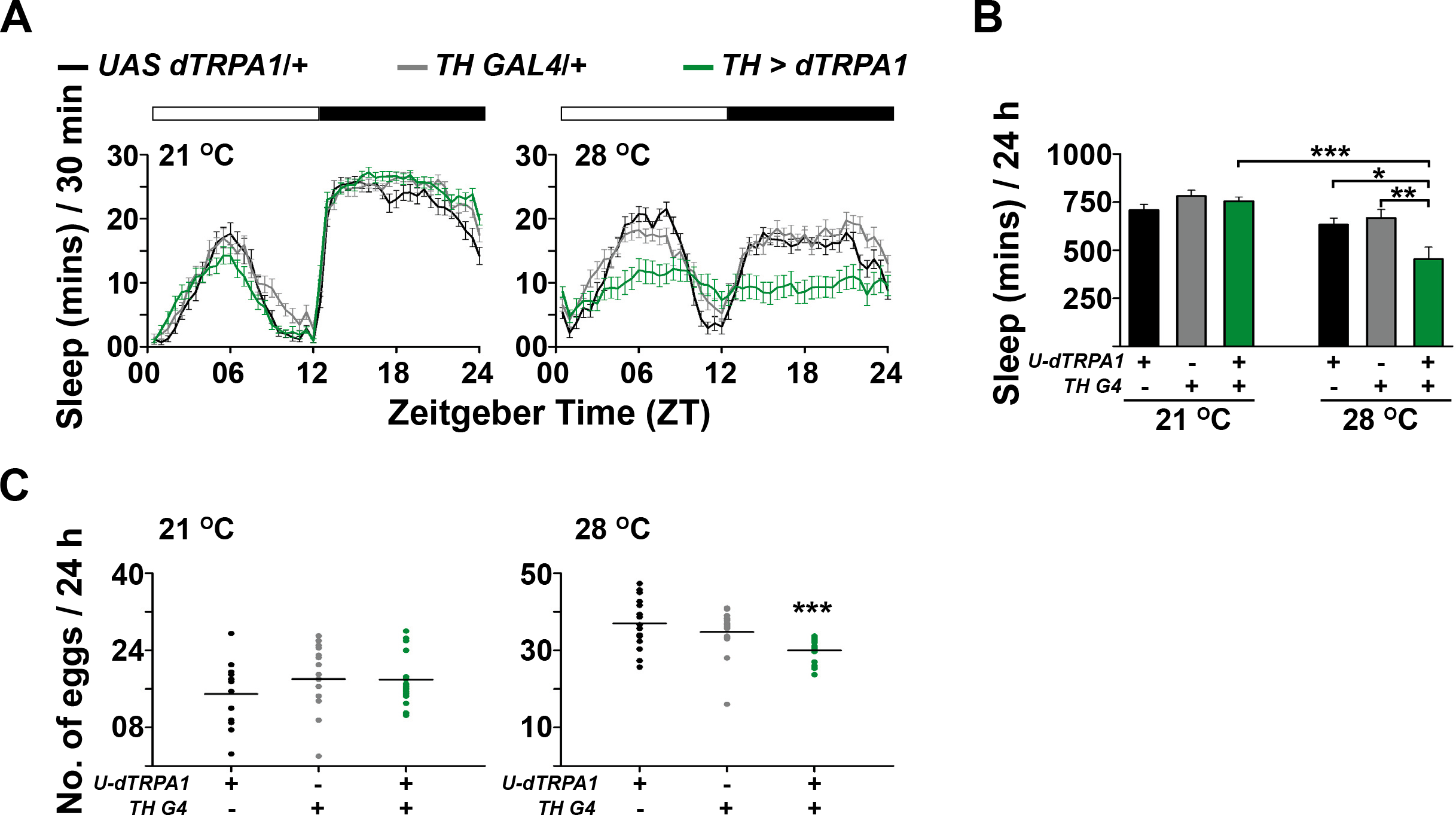
Decreasing sleep levels using dTRPA1-based reversible activation of dopaminergic neurons reversibly decreases egg output (A) Sleep in minutes for every half hour over a period of 24 h averaged across two days at 21 °C (left) and three days at 28 °C (right) is shown for *UAS dTRPA1/+(n* = 29), *TH GAL4/+ (n* = 28) and *TH GAL4* > *UAS dTRPA1 (n* = 32) flies. (B) At 21 °C, total sleep levels of all three genotypes is similar, whereas at 28 °C, *TH GAL4* > *UAS dTRPA1* flies sleep significantly lower than *UAS dTRPA1/+* and *TH GAL4/+* flies (two-way ANOVA with genotype and temperature as fixed factors followed by post-hoc Tukey’s HSD test). (C) Total number of eggs laid averaged across two days at 21 °C (left) is similar across all genotypes, while average number of eggs laid by *TH GAL4* > *UAS dTRPA1 (n* = 16) flies is significantly lower than *UAS dTRPA1/+ (n* = 16) and *TH GAL4/+ (n* = 19) flies during the three days at 28 °C (right, Kruskal-Wallis test). All other details as in Figure 1.

### Transient sleep reduction is accompanied by transient reduction in egg output

It is possible that both caffeine feeding and mechanical perturbation could have broad effects on general physiology of the fly. Therefore, we used a third genetic method whereby sleep reduction is transient and measured egg output following neural-circuit-driven sleep loss. We used the GAL4-UAS system to express a temperature-sensitive cation channel *Drosophila* Transient Receptor Potential 1 *[dTRPA1,* which opens above temperatures of 27 °C and causes hyper-excitation (Hamada et al., 2008)], in dopaminergic neurons that have previously been shown to be wake-promoting (Liu, Liu, Kodama, Driscoll, & Wu, 2012; Shang et al., 2011; Ueno et al., 2012). We recorded sleep levels of flies in tubes and egg output in vials exposed to the following regime - two days at 21 °C followed by three days at 28 °C followed by a day at 21 °C under LD 12:12. As expected, at the higher temperature, sleep was reduced both during daytime and night when dopaminergic neurons were activated, whereas the baseline sleep levels of these experimental flies were not different from that of the parental controls at the lower temperature (Fig 3A-B,). The number of eggs laid by the experimental flies was significantly lower than that of the controls (Fig 3C). Indeed, these differences in egg output between experimental and control flies were not seen at the lower temperature of 21 °C (Fig 3C) when sleep levels were not affected (Fig 3A-B,), suggesting that transiently reducing sleep levels by activating wake-promoting neurons also resulted in transient reduction of egg output. Taken together, our results suggest that sleep loss leads to reduction in egg output, irrespective of the method of sleep deprivation.

### Dopamine transporter mutants show reduced sleep but not reduced egg output in response to caffeine

Given that increasing dopaminergic activity increases wakefulness and decreases egg output, we asked if increasing the amount of dopamine in synaptic clefts also led to decreased egg output. We used flies with loss-of-function mutation in the *fumin (fmn)* gene, which codes for dopamine transporter. Mutant *fmn* flies have been reported to show overall reduced sleep and no reduction in lifespan, but the authors did not measure fertility in their study (Kume, Kume, Park, Hirsh, & Jackson, 2005). We quantified their egg output along with sleep levels and found that the *fmn* flies expectedly showed reduced sleep levels both during the day and night (Fig 4A-B,-top), and the egg output of*fmn* flies was drastically reduced as compared to that of the background control flies (*fmn-bg*, Fig 4C). Once again, we find that flies that sleep less also have low egg output.

**Figure 4.**
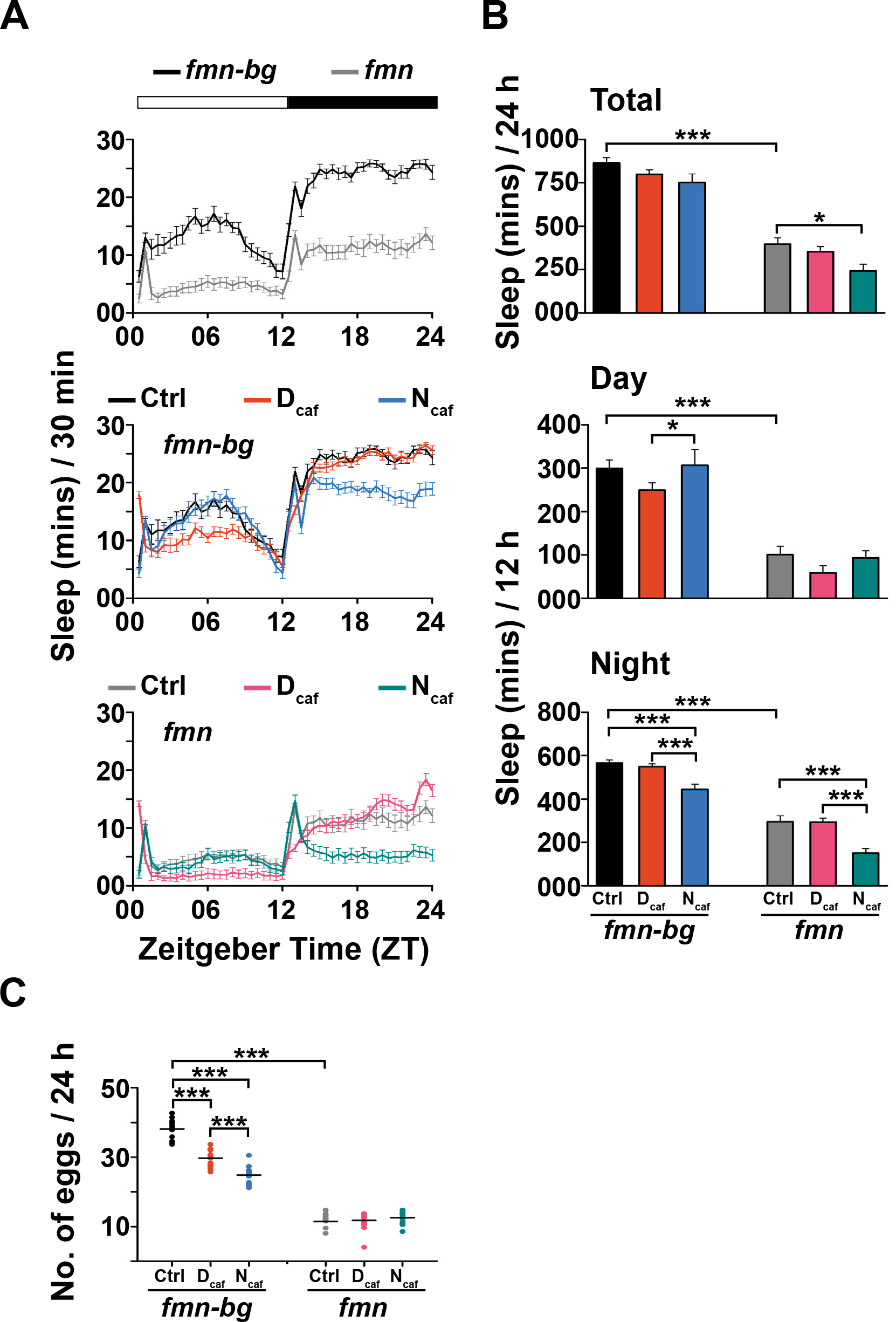
*fmn* flies reduce sleep but not egg output in response to caffeine (A) Sleep in minutes for every half hour over a period of 24 h averaged across 6 days of *fmn* and *fmn* background control *(fmn-bg)* flies (top), *fmn-bg* flies fed with standard food (*n* = 17), caffeine food (0.5 mg/ml) only during the day (D_caf_, *n* = 28) and only during the night (N_caf_, *n* = 26) (middle) and *fmn* receiving control (n = 22), D_caf_ (n = 24) and N_caf_ (n = 28) treatments (bottom). (B) Total sleep levels of *fmn-bg* and *fmn* flies, compared with that of D_caf_ and N_caf_ flies of each genotype (top), daytime sleep (middle) and night sleep (bottom). *fmn* flies sleep significantly lower than the *fmn-bg* flies both during the day and night, thereby leading to overall reduced levels of sleep. Daytime sleep of Dcaf and Ncaf flies of the control genotype are significantly different from one another, whereas night sleep of Ncaf flies is significantly lower than D_caf_ and control flies of the *fmn-bg* genotype. Night sleep of N_caf_ flies is significantly lower than both control and D_caf_ flies of the *fmn* genotype (two-way ANOVA with genotype and treatment as fixed factors followed by post-hoc Tukey’s HSD test). (C) Total number of eggs laid averaged over 6 days by *fmn* flies is significantly lower than that of *fmn-bg* flies (Students’ two-tailed *t* test). D_caf_ flies of*fmn-bg* genotype (n = 14) laid significantly lower number of eggs as compared to its controls (n = 14), while N_caf_ flies of *fmn-bg* genotype (n = 16) laid significantly lower number of eggs as compared to both control and D_caf_ flies. Control (n = 15), D_caf_ (n = 17) and N_caf_ (n = 17) flies of the *fmn* genotype laid similar number of eggs (two-way ANOVA with genotype and treatment as fixed factors followed by post-hoc Tukey’s HSD test). All other details as in Figure 1.

A previous study has shown that *fmn* mutants show a further reduction in sleep when fed with caffeine (Andretic, et al., 2008). We asked if the egg output is also further reduced in *fmn* flies fed with caffeine compared to those fed with standard food. We fed *fmn* and *fmn-bg* flies with 0.5 mg/ml caffeine either only during the day or night and found that N_caf_ flies of both *fmn* and *fmn-bg* genotypes show reduced levels of night sleep as compared to their respective controls (Fig 4B, night), whereas D_caf_ flies of both genotypes show reduced levels of daytime sleep (Fig 4B, day), even though it does not reach statistical significance. Interestingly, just like the previously used inbred flies of the w^1118^ genotype, the *fmn-bg* which are flies from another inbred line show a statistically significant trend of decreasing number of eggs laid by Ctrl, Dcaf and Ncaf flies, in that order (Fig 4C). However, surprisingly, flies of the *fmn* genotype receiving the Ctrl, D_caf_ or N_ca_f treatments did not differ in the average number of eggs laid (Fig 4C). This suggests that while sleep is affected by caffeine treatment in *fmn* flies, egg output is not, suggesting that egg output cannot be reduced by caffeine beyond a threshold. Alternatively, the *fmn* gene may be involved in caffeine-mediated egg output reduction independent of the caffeine-mediated sleep loss.

## Discussion

Our study aimed to understand how sleep affects reproductive output in female fruit flies *Drosophila melanogaster*. We find that feeding flies with caffeine such that it reduces sleep also reduces egg output in both inbred and outbred strains of flies (Figs 1, 2). Moreover, depriving flies of sleep via mechanical perturbation also reduces egg output considerably (Figs 1, 2). A loss-of-function mutation in dopamine transporter gene that results in reduced sleep (Kume, et al., 2005) also results in reduced egg output (Fig 4). Most importantly, reducing sleep by transient dopaminergic neuronal activation reduces egg output; removal of this activation results in wild type levels of sleep and egg output (Fig 3). Thus, these results strongly indicate that it is sleep loss which has a direct detrimental impact on reproductive output. While it is possible that three distinct methods of sleep deprivation all cause a direct negative impact on egg output independent of sleep loss, we feel that it is unlikely, especially considering the transient nature of the genetic manipulation induced sleep loss. To our knowledge, this is the first study to establish a direct link between sleep and reproductive physiology in *Drosophila melanogaster*.

Reproduction in *Drosophila* is regulated by an array of hormones and fecundity critically depends upon balance in the amounts of Juvenile Hormone (JH) and ecdysone (20E) (Soller, Bownes, & Kubli, 1999). Dopamine regulates levels of JH in *Drosophila viridis* (Rauschenbach et al., 2007) thereby indirectly affecting fecundity. Indeed, dopaminergic neuronal circuits are involved in governing oviposition choice, specifically to media containing favourable levels of alcohol (Azanchi, Kaun, & Heberlein, 2013). Moreover, it has been also shown that dopamine acts to promote adaptation of *Drosophila sechelia* to a specialist diet of an otherwise toxic fruit, *Morinda citrifolia* by boosting its fecundity (Lavista-Llanos et al., 2014). In a recent study using genome-wide association methods, two genes encoding dopamine receptors *(Dop1R1* and *DopEcR)* in *D. melanogaster* were shown to have pleiotrophic effects on traits associated with ovariole number and sleep parameters (Lobell, Kaspari, Serrano Negron, & Harbison, 2017). Importantly, lowered levels of dopamine during larval stages or immediately after eclosion both have far reaching consequences in terms of decreased egg output and stalled ovarian development respectively (Neckameyer, 1996). In contrast, we show that a loss-of-function mutation in the dopamine transporter gene which retains dopamine in synaptic clefts reduces sleep and reduces egg output while transient *increase* in dopaminergic activity causes a transient decrease in both sleep and egg output ((Fig 3). Together these results demonstrate that levels of neuromodulatory substances can have strong dose dependent effects such that both low and high titres can lead to sub-optimal outcomes to the organism (Berridge & Arnsten, 2013).

Caffeine is one of the most widely used psychostimulants in the world and it promotes wakefulness and causes sleep deprivation. With increased precedence in shift work and a general lifestyle favouring delayed bedtimes and decreased night sleep levels, the consumption of caffeine specifically during the night is bound to increase. Here, we show that caffeine consumption and increased night activity decreases sleep and negatively alters egg output in *Drosophila.* While we have shown this effect with female flies, it is not wrong to expect similar trends in male reproductive output as well. In conclusion, our results unequivocally show that each method of sleep deprivation, be it chemical, mechanical or genetic, results in sleep loss accompanied with reduction in egg output. For animals that invest in parental care, sleep deprivation may be an inevitable consequence resulting in lowered reproductive output thereby potentially giving rise to a subtle level of parent-offspring conflict or coadaptation. We conclude that sleep may contribute to reproductive success of organisms, thereby amplifying its propensity to be selected for, over evolutionary timescales.

## Materials and Methods

### Fly strains

Fly strains used for both activity/rest and egg output assays were w^1118^ (Bloomington stock # 5905), *fumin (fmn), 2202CS* (background control for *fmn* flies, henceforth referred to as *fmn-bg), TH GAL4, UAS dTRPA1* and previously described outbreeding population Chrono Control Merged [CCM, (Gogna, et al., 2015)]. *Fmn* and *fmn-bg* flies were gifts from Dr. Kazuhiko Kume, Nagoya city University, Nagoya, Japan. Other fly lines were obtained from the Bloomington stock centre, Bloomington, Indiana. All the transgenic flies used were back-crossed into the standard w^1118^ background for at least 7 generations.

### Activity/rest and egg output assays

For the activity/rest assays, 4-5 day old virgin female flies were initially allowed to mate for a day and then were individually housed in tubes (65 mm length, 3 mm diameter) with standard cornmeal food on one end and cotton plug on the other and activity was recorded in DAM2 monitors *(Drosophila* activity monitoring system, Trikinetics, Waltham, Massachusetts, USA). The DAM system works on the standard beam-breaking principle where a fly cuts an infra-red beam whenever it moves in the middle portion of the tube, thereby generating activity counts. Activity counts were binned at 1 min intervals to obtain sleep parameters using the software PySolo (Gilestro & Cirelli, 2009). Flies were housed in light and temperature controlled environments with 12 hours of light and 12 hours of darkness (LD 12:12) at 25 °C using incubators (MIR-273, Sanyo, Japan; DR-36VLC8 Percival Scientific Inc., USA). Flies were flipped into tubes containing either standard food or food containing different concentrations of caffeine (Hi-Media) every 12 hours depending upon their treatment. The activity recording assays were run for a period of 6-7 days. First two days represent baseline days of recording, next three days (days 3-5) were the days during which sleep deprivation was given either by caffeine treatment or temperature increase, and the last two days represent the recovery days during which sleep rebound is expected to occur. For specific assays, flies were fed with caffeine either during day or night for a period of 6 days.

The egg output assays were conducted simultaneously along with the activity/rest assays, on a parallel set of flies housed in glass vials (10 cm length, 2.5 cm diameter) containing ∼3 ml of cornmeal food with or without caffeine depending upon the treatment. For the egg output assays, a small amount of charcoal (0.8 g/L) was added to cornmeal food to increase the contrast between eggs and food surface, thereby aiding in egg counting. As before, flies were transferred into fresh food every 12 h and the number of eggs laid were counted with the help of a stereo-microscope (Olympus, SZ160). In the experiment for sleep deprivation by mechanical means, individual flies were housed in tubes (65 mm in length, 5 mm in diameter) placed in DAM5 monitors which were then mounted on a vortexer (VWR) that was used to mechanically disturb flies either during the day or night. Eggs laid by flies in these tubes as well as by flies that remained undisturbed throughout day or night were then counted for a period of 5 days. Oviposition choice assays were performed by introducing 5 female *w*^1118^ flies for a period of two hours on petri-dishes that contained standard cornmeal food on one half and cornmeal food with specific concentrations of caffeine on the other.

### Statistical analysis

Oviposition preference for a given food was defined as the percentage of total eggs laid on that food surface. Percentage sleep loss was calculated as percentage decrease in sleep during sleep deprivation days with reference to, sleep levels during baseline days. Sleep measures of control and sleep deprived flies were compared using one-way ANOVA with treatment or genotype as a fixed factor followed by post-hoc Tukey’s Honest Significant Difference (HSD) test with p-level set at 0.05. Egg output data were first tested for normality using a Shapiro-Wilk’s W test. One-way ANOVA followed by post-hoc Tukey’s HSD test was conducted if all datasets under consideration were normally distributed. However, even if one of the datasets were not normally distributed, a Kruskal-Wallis test was conducted with p-level set at 0.05.

## Acknowledgments

We thank Todd C. Holmes and Charlotte Helfrich-Forster for helpful comments during writing of the manuscript. We are grateful to Shambhavi Chidambaram for help with validation of some key results. We thank Viveka Singh for help with experiments and useful comments on the manuscript. We thank Rajanna and Muniraju for technical assistance. This work was supported by a research grant (SB/SO/AS/019/2013) awarded by Science and Engineering Research Board, Department of Science and Technology, India.

---

**Supplementary Figure 1.**
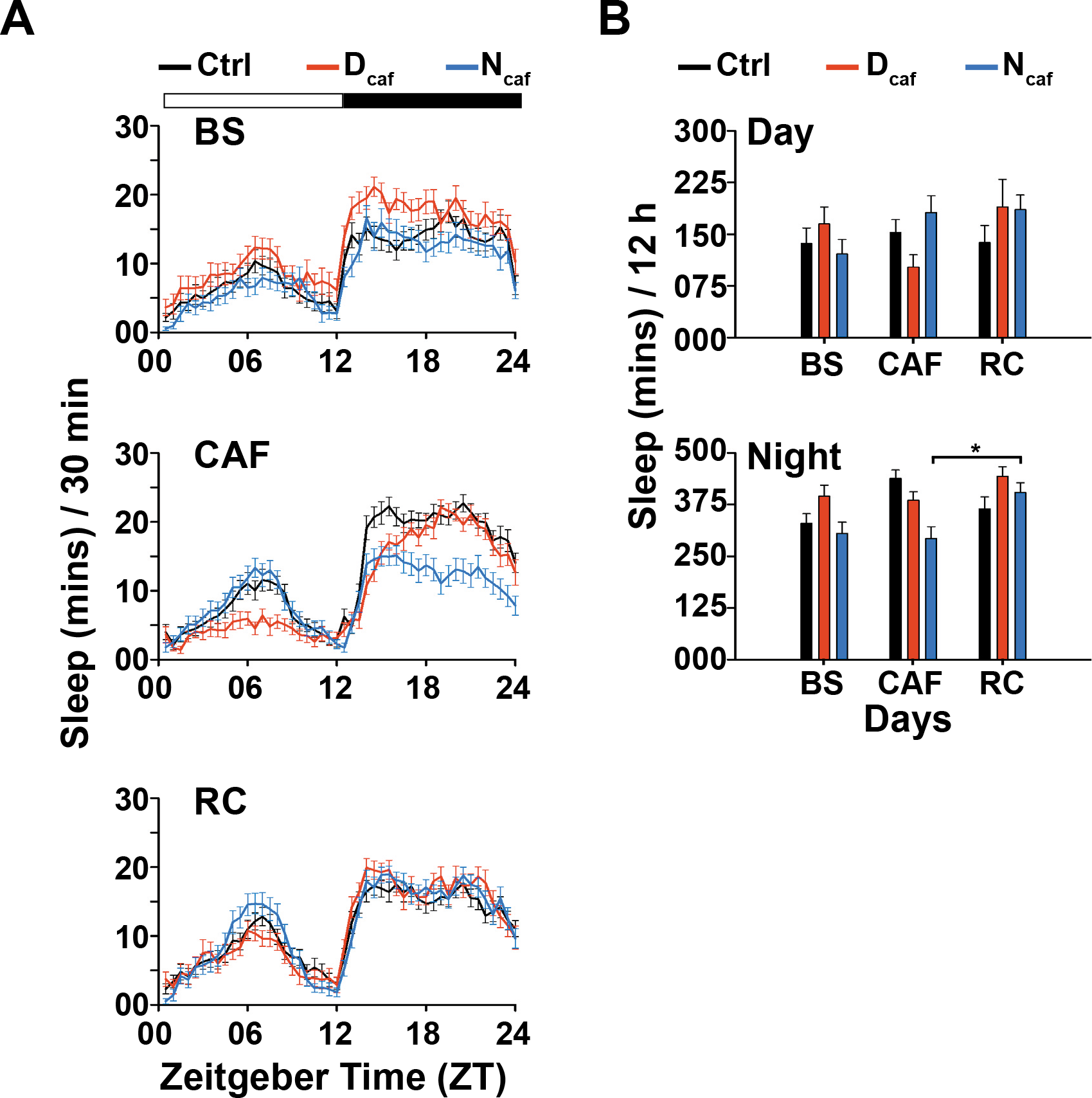
(A) Sleep in minutes for every half hour over a period of 24 h is shown for w^1118^ flies fed with standard food (Ctrl, *n* = 28), flies fed with 1.0 mg/ml caffeine only during the day (D_ca_f, *n* = 29) and only during the night (N_caf_, *n* = 28) averaged across two baseline (BS), three caffeine feeding (CAF) and two recovery (RC) days. (B) Daytime (top) and night (bottom) sleep of control, D_caf_ and N_caf_ flies are compared across BS, CAF and RC days. Only night sleep of N_caf_ flies during CAF and RC days is significantly different from each other (two-way ANOVA with treatment and days as fixed factors followed by post-hoc Tukey’s HSD test). All other details as in Figure 1.

**Supplementary Figure 2.**
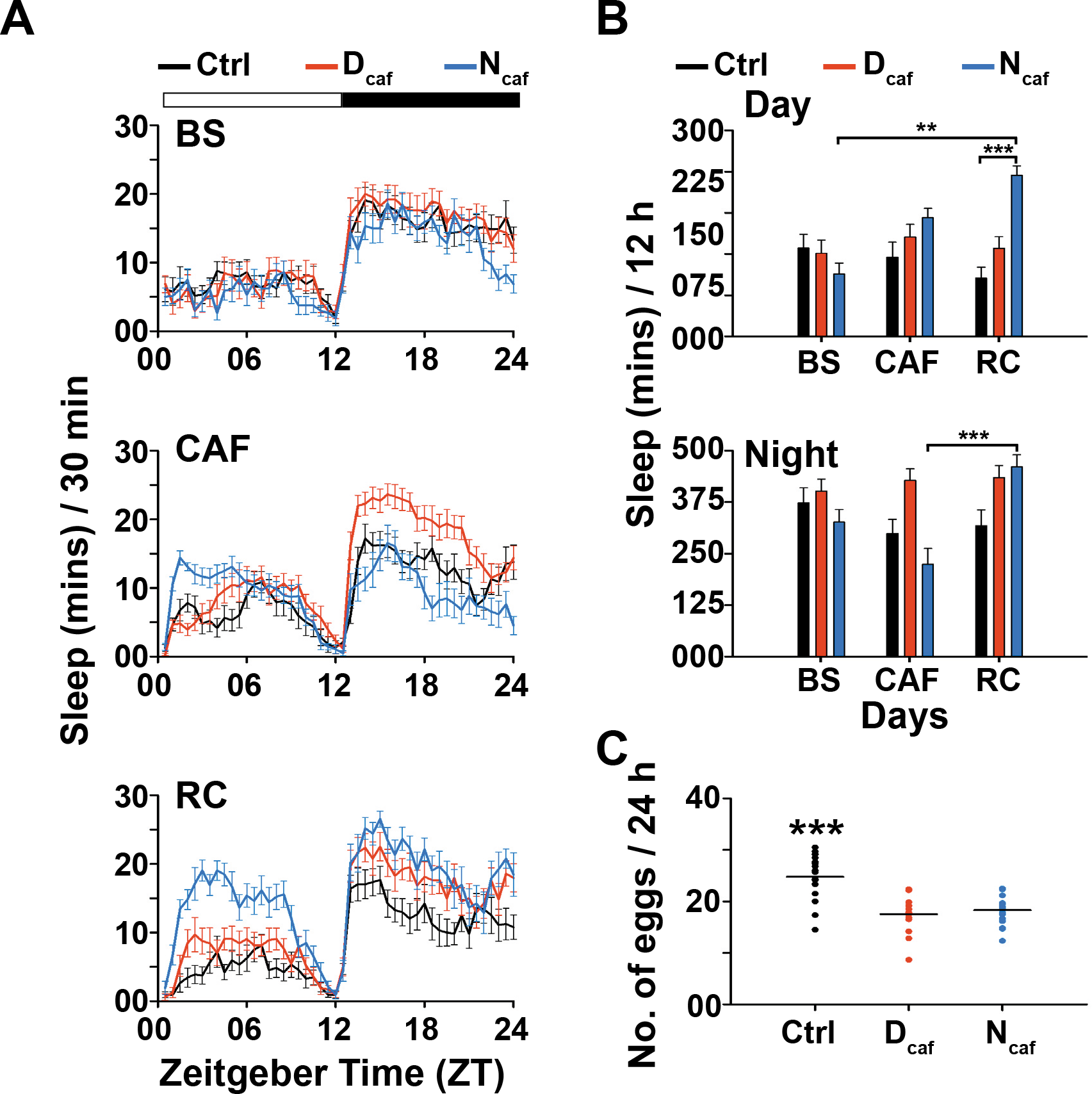
(A) Sleep in minutes for every half hour over a period of 24 h is shown for control flies of outbred CCM population fed with standard food (Ctrl, *n* = 16), flies fed with caffeine only during the day (D_caf_, *n =* 16) and only during the night (N_ca_f, *n =* 14) for caffeine concentration of 4.0 mg/ml averaged across two baseline (BS), three caffeine feeding (CAF) and two recovery (RC) days. Night sleep of N_caf_ flies during CAF days is lower than that of controls, and both daytime and night sleep of N_caf_ flies is higher than the controls during RC. (B) Daytime sleep levels of control and Dcaf flies show no differences across different days, whereas those of control and Ncaf flies significantly differ from each other during RC. Daytime sleep of N_caf_ flies during RC is significantly higher than that during BS. Night sleep of Ncaf flies during CAF and RC days are significantly different from each other other (two-way ANOVA with treatment and days as fixed factors followed by post-hoc Tukey’s HSD test). (C) Total eggs laid by control *(n* = 16), D_caf_ (n = 14) and N_caf_ (n = 18) flies averaged across six days of caffeine feeding. Control flies laid higher number of eggs as compared to both D_caf_ and N_caf_ flies (Kruskal-Wallis test). All other details as in Figure 1.

**Supplementary Figure 3.**
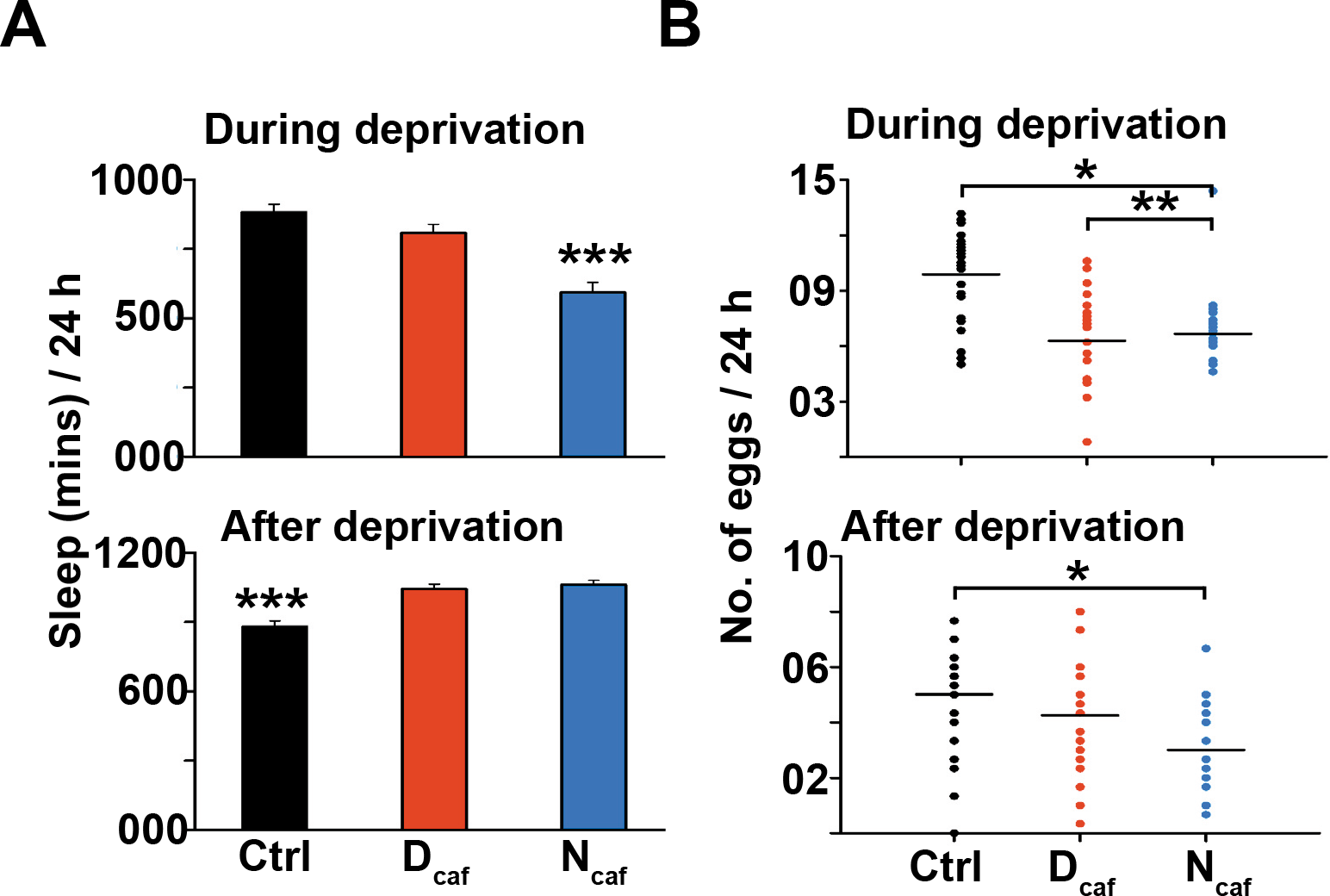
(A) Total sleep (top) during 6 days of sleep deprivation and (bottom) averaged for 3 days post-deprivation. Sleep of N_dep_ (n = 16) flies is significantly lower than both control (n = 29) and D_dep_ (n = 21) flies during sleep deprivation, whereas both D_dep_ and N_dep_ flies sleep more after deprivation (one-way ANOVA followed by post-hoc Tukey’s HSD test). (B) Average number of eggs laid (top) during sleep deprivation and (bottom) after sleep deprivation. D_dep_ and N_dep_ flies lay lesser number of eggs as compared to control flies during deprivation, but only Ndep flies lay lower number of eggs compared to control flies after deprivation (Kruskal-Wallis tests). All other details as in Figure 1.

## REFERENCES

Alvarenga, T. A., Hirotsu, C., Mazaro-Costa, R., Tufik, S., & Andersen, M. L. (2015). Impairment of male reproductive function after sleep deprivation. Fertil Steril, 103(5), 1355–1362 e1351.

Andersen, M. L., & Tufik, S. (2002). Distinct effects of paradoxical sleep deprivation and cocaine administration on sexual behavior in male rats. Addict Biol, 7(2), 251–253.

Andretic, R., Kim, Y. C., Jones, F. S., Han, K. A., & Greenspan, R. J. (2008). Drosophila D1 dopamine receptor mediates caffeine-induced arousal. Proc Natl Acad Sci U S A, 105(51), 20392–20397.

Azanchi, R., Kaun, K. R., & Heberlein, U. (2013). Competing dopamine neurons drive oviposition choice for ethanol in Drosophila. Proc Natl Acad Sci US A, 110(52), 21153–21158.

Beaver, L. M., Gvakharia, B. O., Vollintine, T. S., Hege, D. M., Stanewsky, R., & Giebultowicz, J. M. (2002). Loss of circadian clock function decreases reproductive fitness in males of Drosophila melanogaster. Proc Natl Acad Sci U S A 99(4), 2134–2139.

Berridge, C. W., & Arnsten, A. F. (2013). Psychostimulants and motivated behavior: arousal and cognition. Neurosci Biobehav Rev, 37(9 Pt A), 1976–1984.

Bushey, D., Hughes, K. A., Tononi, G., & Cirelli, C. (2010). Sleep, aging, and lifespan in Drosophila. BMC Neurosci, 11, 56.

Cirelli, C., Bushey, D., Hill, S., Huber, R., Kreber, R., Ganetzky, B., et al.. (2005). Reduced sleep in Drosophila Shaker mutants. Nature, 434(7037), 1087–1092.

Everson, C. A., Bergmann, B. M., & Rechtschaffen, A. (1989). Sleep deprivation in the rat: III. Total sleep deprivation. Sleep, 12(1), 13–21.

Everson, C. A., & Szabo, A. (2009). Recurrent restriction of sleep and inadequate recuperation induce both adaptive changes and pathological outcomes. Am J Physiol Regul Integr Comp Physiol, 297(5), R1430–1440.

Everson, C. A., & Szabo, A. (2011). Repeated exposure to severely limited sleep results in distinctive and persistent physiological imbalances in rats. PLoS One, 5(8), e22987.

Gilestro, G. F., & Cirelli, C. (2009). pySolo: a complete suite for sleep analysis in Drosophila. Bioinformatics, 25(11), 1466–1467.

Gogna, N., Singh, V. J., Sheeba, V., & Dorai, K. (2015). NMR-based investigation of the Drosophila melanogaster metabolome under the influence of daily cycles of light and temperature. Mol Biosyst, 11(12), 3305–3315.

Hamada, F. N., Rosenzweig, M., Kang, K., Pulver, S. R., Ghezzi, A., Jegla, T. J., et al.. (2008). An internal thermal sensor controlling temperature preference in Drosophila. Nature, 454(7201), 217–220.

Hendricks, J. C., Finn, S. M., Panckeri, K. A., Chavkin, J., Williams, J. A., Sehgal, A., et al.. (2000). Rest in Drosophila is a sleep-like state. Neuron, 25(1), 129–138.

Kayser, M. S., Mainwaring, B., Yue, Z., & Sehgal, A. (2015). Sleep deprivation suppresses aggression in Drosophila. Elife, 4, e07643.

Kloss, J. D., Perlis, M. L., Zamzow, J. A., Culnan, E. J., & Gracia, C. R. (2015). Sleep, sleep disturbance, and fertility in women. Sleep Med Rev, 22, 78–87.

Krause, A. J., Simon, E. B., Mander, B. A., Greer, S. M., Saletin, J. M., Goldstein-Piekarski, A. N., et al.. (2017). The sleep-deprived human brain. Nat Rev Neurosci, 18(7), 404–418.

Kume, K., Kume, S., Park, S. K., Hirsh, J., & Jackson, F. R. (2005). Dopamine is a regulator of arousal in the fruit fly. J Neurosci, 25(32), 7377–7384.

Lavista-Llanos, S., Svatos, A., Kai, M., Riemensperger, T., Birman, S., Stensmyr, M. C., et al.. (2014). Dopamine drives Drosophila sechellia adaptation to its toxic host. Elife, 3.

Liu, Q., Liu, S., Kodama, L., Driscoll, M. R., & Wu, M. N. (2012). Two dopaminergic neurons signal to the dorsal fan-shaped body to promote wakefulness in Drosophila. Curr Biol, 22(22), 2114–2123.

Lobell, A. S., Kaspari, R. R., Serrano Negron, Y. L., & Harbison, S. T. (2017). The Genetic Architecture of Ovariole Number in Drosophila melanogaster: Genes with Major, Quantitative, and Pleiotropic Effects. G3 (Bethesda).

Loh, D. H., Kuljis, D. A., Azuma, L., Wu, Y., Truong, D., Wang, H. B., et al.. (2014). Disrupted reproduction, estrous cycle, and circadian rhythms in female mice deficient in vasoactive intestinal peptide. J Biol Rhythms, 29(5), 355–369.

Mahoney, M. M. (2010). Shift work, jet lag, and female reproduction. Int J Endocrinol, 2010, 813764.

Neckameyer, W. S. (1996). Multiple roles for dopamine in Drosophila development. Dev Biol, 176(2), 209–219.

Nitz, D. A., van Swinderen, B., Tononi, G., & Greenspan, R. J. (2002). Electrophysiological correlates of rest and activity in Drosophila melanogaster. Curr Biol, 12(22), 1934–1940.

Rauschenbach, I. Y., Chentsova, N. A., Alekseev, A. A., Gruntenko, N. E., Adonyeva, N. V., Karpova, E. K., et al.. (2007). Dopamine and octopamine regulate 20-hydroxyecdysone level in vivo in Drosophila. Arch Insect Biochem Physiol, 65(2), 95–102.

Rechtschaffen, A., Gilliland, M. A., Bergmann, B. M., & Winter, J. B. (1983). Physiological correlates of prolonged sleep deprivation in rats. Science, 221(4606), 182–184.

Seugnet, L., Suzuki, Y., Thimgan, M., Donlea, J., Gimbel, S. I., Gottschalk, L., et al.. (2009). Identifying sleep regulatory genes using a Drosophila model of insomnia. J Neurosci, 29(22), 7148–7157.

Shang, Y., Haynes, P., Pirez, N., Harrington, K. I., Guo, F., Pollack, J., et al.. (2011). Imaging analysis of clock neurons reveals light buffers the wake-promoting effect of dopamine. Nat Neurosci, 14(7), 889–895.

Shaw, P. J., Cirelli, C., Greenspan, R. J., & Tononi, G. (2000). Correlates of sleep and waking in Drosophila melanogaster. Science, 287(5459), 1834–1837.

Shaw, P. J., Tononi, G., Greenspan, R. J., & Robinson, D. F. (2002). Stress response genes protect against lethal effects of sleep deprivation in Drosophila. Nature, 417(6886), 287–291.

Sheeba, V., Sharma, V. K., Shubha, K., Chandrashekaran, M. K., & Joshi, A. (2000). The effect of different light regimes on adult life span in Drosophila melanogaster is partly mediated through reproductive output. J Biol Rhythms, 15(5), 380–392.

Soller, M., Bownes, M., & Kubli, E. (1999). Control of oocyte maturation in sexually mature Drosophila females. Dev Biol, 208(2), 337–351.

Summa, K. C., Vitaterna, M. H., & Turek, F. W. (2012). Environmental perturbation of the circadian clock disrupts pregnancy in the mouse. PLoS One, 7(5), e37668.

Ueno, T., Tomita, J., Tanimoto, H., Endo, K., Ito, K., Kume, S., et al.. (2012). Identification of a dopamine pathway that regulates sleep and arousal in Drosophila. Nat Neurosci, 25(11), 1516–1523.

van Alphen, B., Yap, M. H., Kirszenblat, L., Kottler, B., & van Swinderen, B. (2013). A dynamic deep sleep stage in Drosophila. J Neurosci, 22(16), 6917–6927.

Wu, M. N., Ho, K., Crocker, A., Yue, Z., Koh, K., & Sehgal, A. (2009). The effects of caffeine on sleep in Drosophila require PKA activity, but not the adenosine receptor. J Neurosci, 29(35), 11029–11037.

